# Differences in neutralization susceptibility between clade C HIV viruses from breastmilk versus contemporaneous circulating viruses from sexually acquired infections

**DOI:** 10.1101/2025.11.04.686466

**Authors:** Elena E. Giorgi, Kevin Gillespie, Elizabeth Domin, Genevieve Fouda, Sallie R. Permar, David C. Montefiori, Holly Janes

**Affiliations:** Vaccine and Infectious Disease Division, Fred Hutchinson Cancer Center, Seattle; Duke Human Vaccine Institute, Duke University School of Medicine, Durham, NC, USA; Weill Cornell Medicine, Department of Pediatrics, Division of Infectious Diseases, New York, New York; Department of Surgery, Duke University School of Medicine, Durham, NC; Department of Biostatistics, University of Washington, Seattle, WA

## Abstract

HIV viruses that establish infection possess phenotypic and genotypic characteristics that have been selected for and that differ across transmission routes, including their susceptibility to broadly neutralizing antibodies (bnAbs). While sexually transmitted viruses have been well characterized, studies of vertically transmitted viruses are sparse and from cohorts that are often small in size and more than a decade old. To investigate whether viruses transmitted vertically during lactation possess distinct neutralization profiles compared to viruses transmitted sexually, we compared the neutralization sensitivity of 25 clade C breastmilk viruses to that of 99 contemporaneous clade C viruses from sera of adults with sexual acquisition against three bnAbs in clinical development.Three out of 7 breastmilk donors (43%) had one or more viruses resistant to 2 or more bnAbs, compared to 8 out of 99 (8%) contemporaneous adult viruses (p=0.02). Breastmilk viruses were more resistant to PGT121 and VRC07.523 (median IC80 >50 compared to 1.16 for PGT121, and 12.75 vs. 0.38 for VRC07.523; p=0.013 and <0.001 respectively), and more breastmilk viruses than adult viruses were resistant to VRC07.523 (94% vs. 43%, p=0.001). Interestingly, the breastmilk viruses most resistant to VRC07.523 had on average one or more glycans in V3 compared to adult transmitted viruses (median 3 vs. 2 glycosylation sites, including flanking position 295; p=0.009), and the number of V3 glycans was negatively correlated with VRC07.523 sensitivity (p=0.007). These findings highlight potential differences in bnAb susceptibility of vertically transmitted viruses and emphasize the need to increase sequencing efforts and screening of infant viruses to better inform the efficacy of candidate bnAbs to prevent vertical transmission of HIV.

## Introduction

The HIV pandemic remains a worldwide health challenge, with ~40 million people living with HIV (PLWH) in 2023, of which 1.4 million (4%) are 14 or younger (1). In 2023, ~120,000 new pediatric HIV infections occurred, representing 10% of all new infections, although newborns represent 1% of the global population (1). Children accounted for 12% of all AIDS-related deaths (2), an estimate likely to increase if current threats to HIV prevention and care remain in place (2).

HIV vertical transmission via breastfeeding occurs through the oral and gastric mucosa from virus that resides and replicates in both cell-free and cellular breastmilk compartments (3). Upon transmission, HIV undergoes a significant bottleneck in genetic diversity (4–7). In adult sexual transmissions, early transmitted viruses have been shown to have distinct phenotypic differences (8–10), and there is evidence that phenotypic selection also occurs in vertical transmission (5, 7, 11–13), albeit from viruses sampled over a decade ago (14). Over time, HIV has accumulated greater global diversity and increased resistance against certain bnAbs (15–22).

Here we evaluate the potency of three bnAbs in clinical development for prevention of both pediatric and adult HIV—PGDM1400.LS, VRC07.523LS, and the IAVI bnAb PGT121.414LS (23, 24) against 25 clade C breastmilk (BM) sequences of the HIV envelope (env) gene sampled from breastfeeding PLWH. We compare the BM env neutralization titers with that of contemporaneous clade C viruses sampled from serum of adults acquiring HIV sexually and identify unique phenotypic and genotypic characteristics of the BM viruses. Our results highlight the importance of HIV sequencing of contemporary vertically transmitted viruses to inform bnAb clinical development.

## Results

### Phylogenetic Analysis

Phylogenetic analysis of the 130 clade C Env sequences shows within-host clustering of the BM sequences (Fig 1). Hierarchical clustering of the IC80 titers shows no preferential clustering of the titers from the BM viruses and infant viruses compared to contemporaneous adult viruses (Fig. 2). However, the cluster with the most resistant viruses (top four rows in Fig. 2) comprised of one infant virus, two BM viruses from a transmitting mother (participant 1209), and one contemporaneous adult virus. All mothers had at least one env with IC80>10 against at least one of the three bnAbs (Fig. 2).

**Fig 1.**
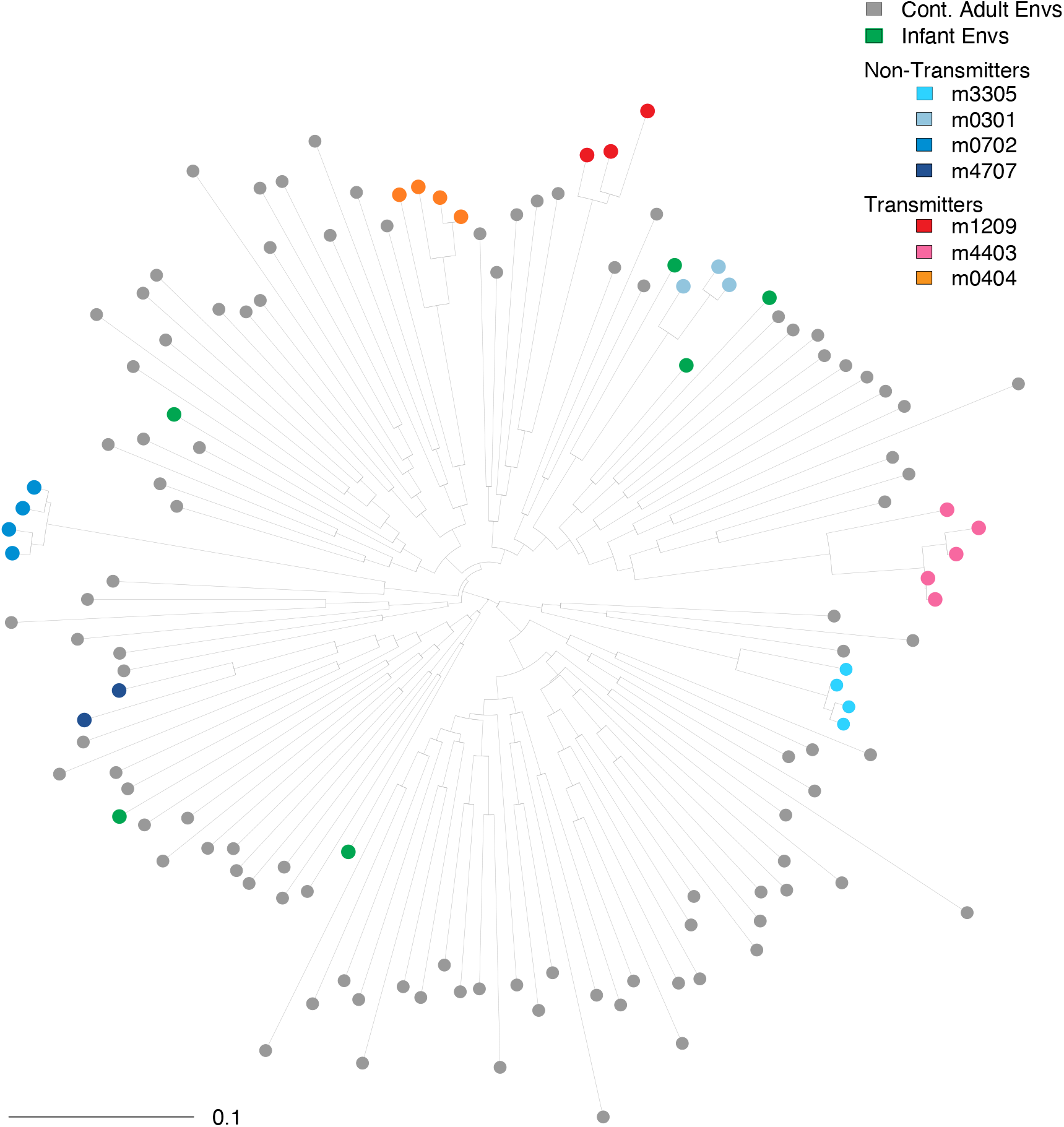
Midpoint rooted phylogenetic tree of 130 clade C Env sequences sampled from BM (N=25), infants (N=6), and from contemporaneous adult transmissions (N=99, gray filled circles). Envs from transmitting mothers are color-coded using warm colors and cold colors are used for Envs from non-transmitting mothers.

**Fig 2.**
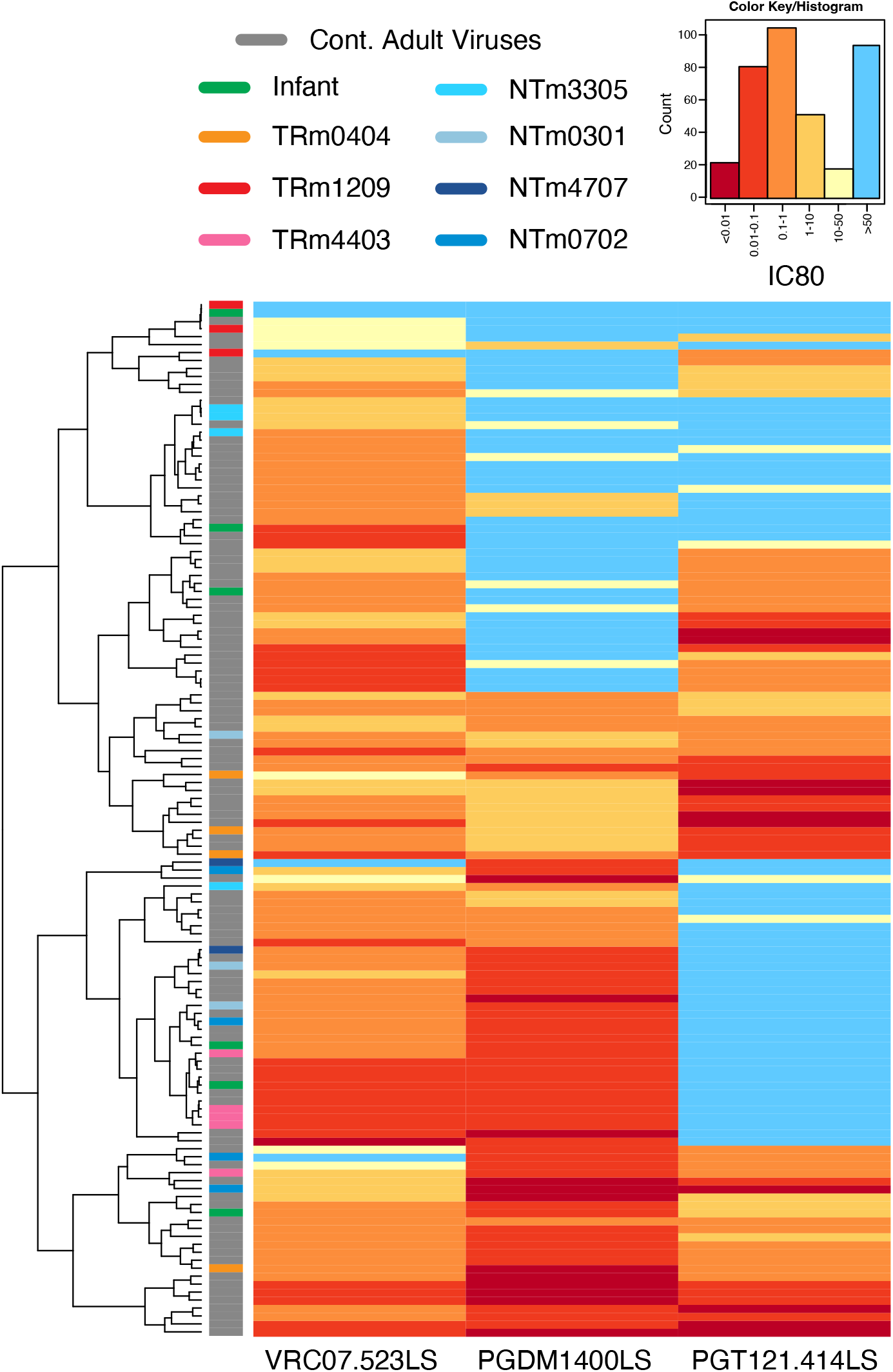
Heatmap visualization of the IC80 titers of all 130 Envs (rows) against the three bnAbs VRC07.523, PGT121, and PGDM1400 (columns). Titers are color-coded with warmer colors indicating stronger susceptibility and lighter colors indicating resistance, with light blue indicating above titers above the detection threshold of 50. Rows are hierarchically clustered and color-coded bars follow the same schema as in Fig. 1: warm colors for transmitting mothers, cold colors for non-transmitting mothers, and gray for contemporaneous adult

### BM viruses were more resistant to VRC07.523LS and PGT121 compared to contemporaneous adult viruses

VRC07.523LS was the most potent, both in breadth and magnitude, of the three bnAbs: it neutralized 117 out of 130 (90%) viruses (IC80≤10 μg per milliliter; Fig. 2 and 3). PGT121 and PGDM1400 neutralized 70 (54%) and 85 (65%) of viruses, respectively.

**Fig 3.**
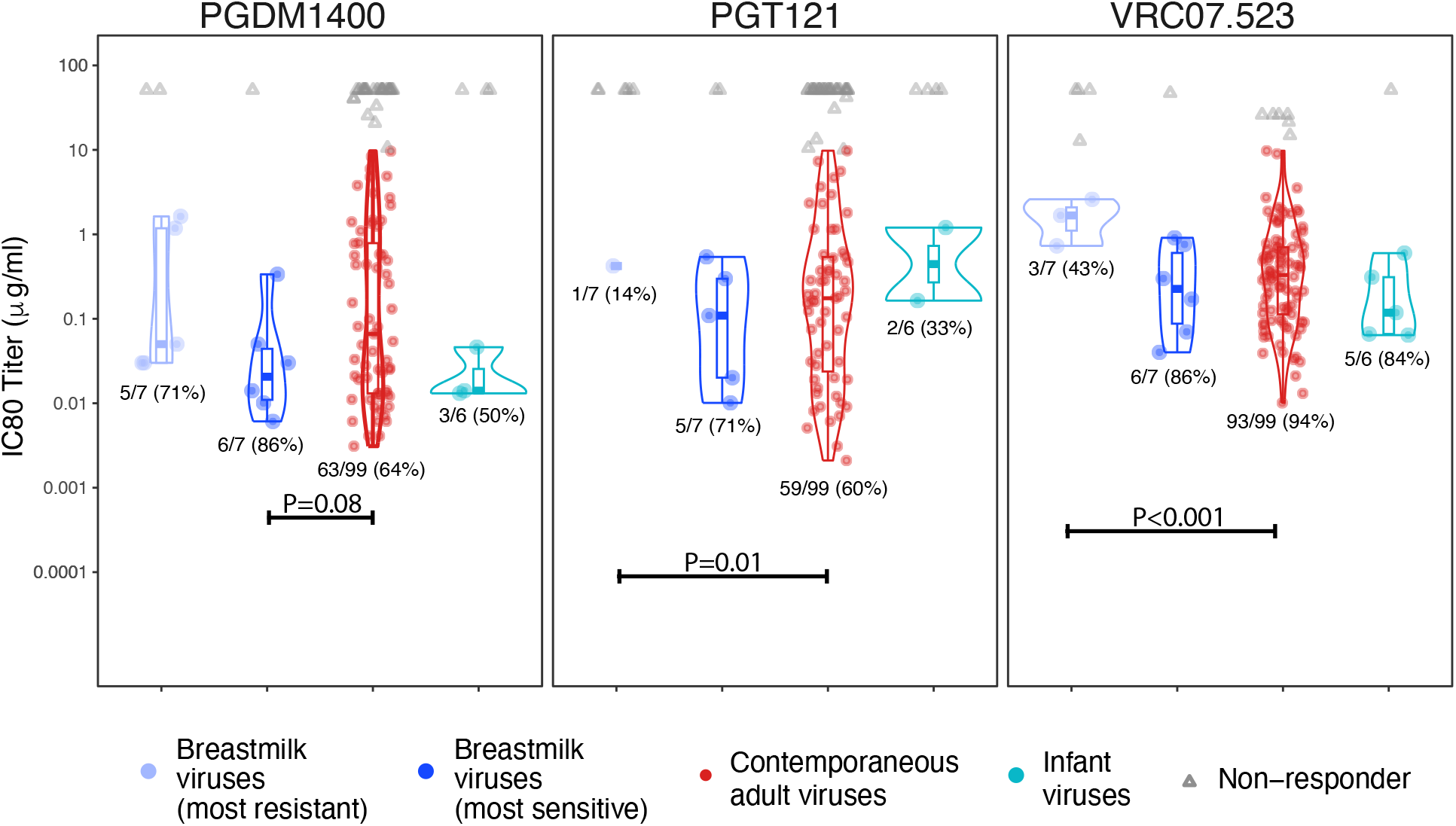
Distribution of IC80 titers against VRC07.523, PGT121, and PGDM1400, for breastmilk viruses, contemporaneous adult viruses, and infant viruses. Two sets of breastmilk viruses are considered: the virus most sensitive to a given bnAb, among all viruses sequenced from each mother; and the virus most resistant to a given bnAb.

Complete resistance (IC80 >50) to all three bnAbs was observed in one infant virus (BF942.218d) and one BM virus (1209.bm_H5), but none of the contemporaneous adult viruses (Fig 2). Among CH009 study participants, three (1209, 3305, and 4707) had at least one virus resistant to two or more bnAbs, whereas only 8 of 99 contemporaneous adult viruses were resistant to two or more bnAbs (p=0.02 by Fisher exact; Fig. 2).

In the most sensitive datasets, BM viruses were marginally more sensitive than contemporaneous clade C viruses to PGDM1400 (p=0.08), however, this was no longer observed in most resistant BM viruses (Fig. 3 and Supp. Fig S3). In the most resistant datasets, BM viruses were more resistant than contemporaneous adult viruses to PGT121 (median IC80 >50 and 1.16 respectively, p=0.01) and VRC07.523LS (median IC80 12.75 vs. 0.38 respectively, p<0.001; Fig. 3). Additionally, more BM viruses than contemporaneous adult viruses were resistant to VRC07.523LS: the response rate was higher in clade C contemporaneous viruses compared to BM viruses (94% vs. 43%, p=0.001). Similar trends were observed for IC50 titers (Supp. Fig. S1 and S2).

### Residues associated with bnAb resistance were more prevalent in BM viruses compared to contemporaneous adult viruses

We identified 96 and 43 sites (see Methods) associated with changes in neutralization susceptibility to PGT121 and VRC07.523LS, respectively, all published in Bricault et al., 2019 (25). A total of 60 sites were considered for PGDM1400, compiled from analyses conducted in Bricault et al., 2019 (25), Roark et al., 2021 (26), and Mishra et al., 2020 (27). After excluding sites where no variation was observed across all sequences, the final analysis included 27, 48, and 24 sites for PGDM1400, PGT121 and VRC07.523LS respectively.

No statistically significant difference in amino acid prevalence between BM and contemporaneous adult viruses was found at signature sites associated with VRC07.523LS neutralization. There were however significant differences in amino acid frequencies at three and two signature sites associated with PGDM1400 and PGT121 respectively (Fig. 4). At each site, the resistance-associated residue was more frequent in BM viruses, whereas the sensitive-associated residue was more frequent in contemporaneous adult viruses.

**Fig 4.**
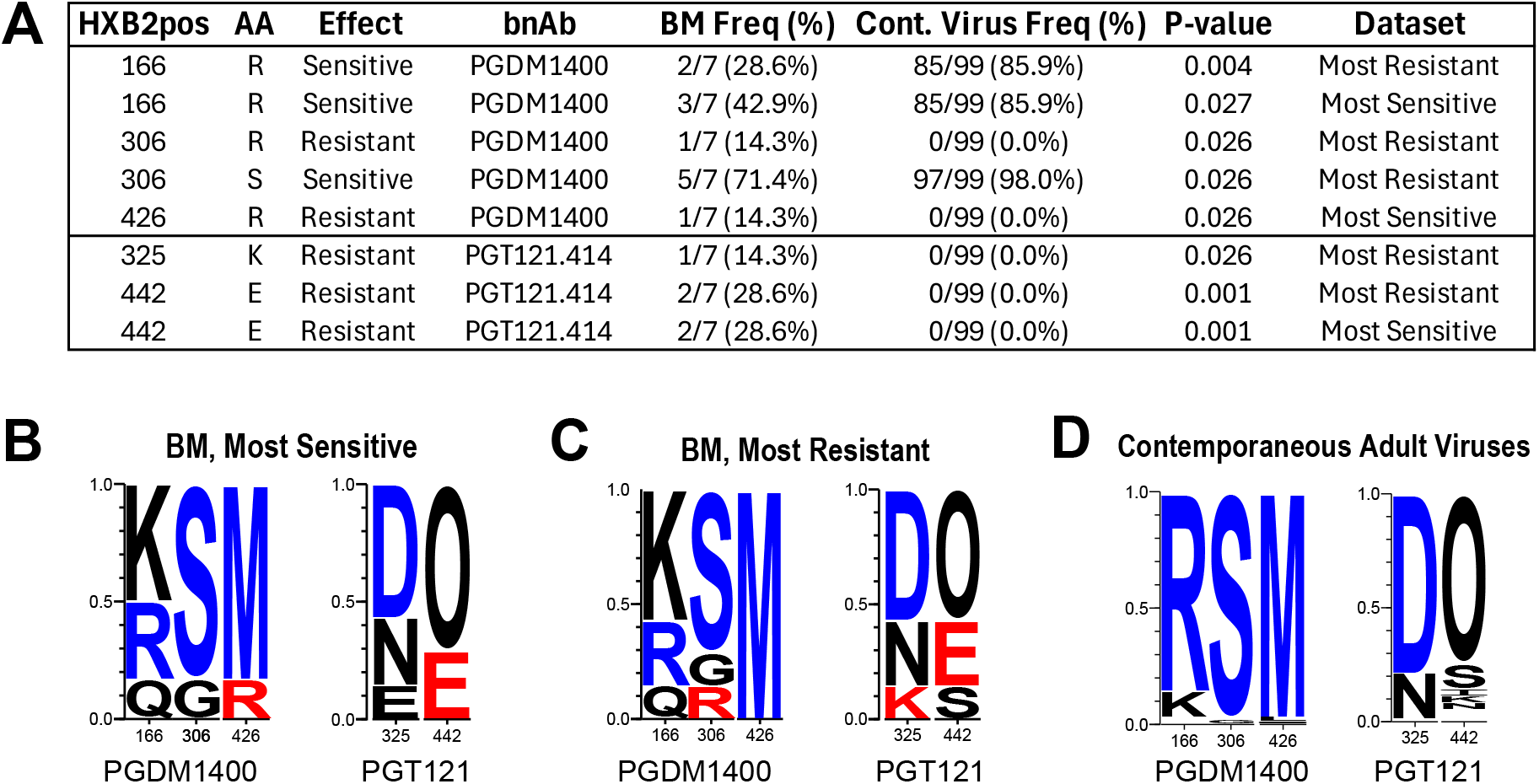
(A) Sites associated with changes in bnAb susceptibility with significantly different prevalence of residue between BM viruses and contemporaneous adult viruses. Signature residues 166R and 306S, in V2 and V3 respectively, are both associated with enhanced sensitivity to PGDM1400 [32], and were both found at a lower frequency in the BM viruses (29% vs. 86% for 166R, p=0.004; 71% vs. 98% for 306S, p=0.03). On the other hand, PGDM1400 resistance-associated signature residues 306R and 426R, and PGT121 resistance-associated signature residue 442E were more prevalent in the BM viruses compared to the adult viruses (14% vs. 0% for both 306R and 426R, p=0.03; 28% vs. 0% for 442E). **(B-D)** Logo plots of sites listed in panel A for each dataset. Each logo is proportional to the frequency of the corresponding amino acid at that site. Logos are color-coded red if found to be associated with increased bnAb resistance, blue if associated with increased sensitivity, and black if neither. O indicates an asparagine (N) embedded in a glycan motif (N followed by any AA except proline, followed by either S or T).

### BM viruses most resistant to VRC07.523LS carry a higher number of glycosylation sites in V3

Env length, electrochemical charge, and number of glycosylation sites in the five variable regions (V1-V5) have also been shown to be associated with changes in bnAb susceptibility (25). On average, BM Envs in the VRC07.523LS most resistant set had one more glycosylation site than contemporaneous adult viruses in or around V3 (median 3 vs. 2 glycosylation sites, including flanking position 295; p = 0.009), with the additional sequon found at either positions 295, 332, or both. Interestingly, in all viruses combined, we observed a significant positive correlation between VRC07.523LS IC80 titers and number of V3 glycosylation sites (τ=0.21, p=0.007; Supp. Fig. S3).

Compared to adult viruses, PGT121 most resistant BM viruses carried a higher number of V2 glycosylation sites (p=0.02), PGDM1400 most resistant BM viruses had a lower V4 electrochemical charge (p=0.02), and PGT121 most sensitive BM viruses carried a higher number of V2 glycosylation sites (p=0.03; Supp. Fig. S4).

## Discussion

We characterized the neutralization susceptibility of 25 clade C viruses sampled from breastmilk from 7 people living with HIV, against VRC07.523LS, PGT121, and PGDM1400, in comparison to 99 clade C viruses sampled from adults who acquired HIV sexually. Breastmilk viruses were significantly more resistant to VRC07.523LS (median IC80 = 12.75 vs. 0.38 for adult viruses). While no difference in residue prevalence was found at VRC07.523LS contact sites, BM viruses resistant to VRC07.523 had a higher number of glycosylation sites in V3 compared to contemporaneous adult viruses. Glycans in V3 have been observed to affect HIV infectivity, tropism, and coreceptor usage (28, 29). Interestingly, autologous responses to V3 have been found to be associated to vertical transmission risk in peripartum infections (30, 31).

With clinical trials under way to test the safety of combination bnAb infusions to prevent HIV acquisition in infants, it is important to compare the neutralization susceptibility of vertically transmitted viruses to those sampled from sexual transmissions. Vertically transmitted viruses, and particularly full env sequences from vertical acute transmissions, are seldom sampled: few are currently available from GenBank, and most are over a decade old (14).

While limited in sample size, the trends described here, together with findings from previous studies (5, 11, 13, 32–34), suggest that distinct transmission routes may select for different phenotypes, including bnAb susceptibility. More infant env sequencing, particularly in acute infections, is needed to ensure that bnAbs selected for clinical development are potent and effective against both horizontally and vertically transmitted viruses.

## Methods

### Study Participants

Twenty-five BM sequences of the HIV envelope gene (env) were obtained from seven mothers in CH009, a Malawian cohort of breastfeeding PLWH and their infants, enrolled at birth between 2008 and 2009 (12, 35). Three mothers transmitted HIV to their infants during follow-up and four did not. The median number of env sequences per participant was 4 (range 2-5; Fig. 1). Six plasma env sequences from unrelated infants from the same cohort who acquired HIV via breastmilk were also included, one env per infant (12). For comparison, we used 99 clade C env sequences obtained from serum of adults from Malawi, Botswana and South Africa who acquired HIV sexually between 2006 and2008 (19). We refer to these as “contemporaneous adult viruses” and to maternal BM-derived envs as “BM viruses.”

### Neutralization Assays

Neutralization titers for all 25 BM viruses and 6 infant viruses against bnAbs VRC07-523LS, PGT121.414LS, PGDM1400LS were assessed using the TZM-bl neutralization assay (36–38). Neutralizing antibodies were measured as a function of reductions in luciferase (Luc) reporter gene expression after a single round of infection in TZM-bl cells as previously described (36–38). Contemporaneous adult viruses were similarly assayed against VRC07-523LS, PGT121, and PGDM1400 (19). PGT121.414LS is an engineered versions of PGT121 engineered for improved manufacturability (39) and is nearly equivalent to non-engineered PGT121 in neutralization potency and breadth (Mike Seaman, personal communication).

Similarly, while BM viruses were measured against the LS versions of PGDM1400, whereas the contemporaneous adult viruses were not, we do not expect the LS mutation to affect neutralization titers (40, 41) (42). For simplicity, we refer to these bnAbs as PGT121 and PGDM1400, with the understanding that the antibodies tested were different versions in the two virus sets.

### Phylogenetic Analyses

Phylogenetic trees were generated from the amino acid alignment using the software IQ Tree (43) with a Gamma model for rate heterogeneity and ultrafast bootstrap branch support. Hierarchical clustered heatmaps of the neutralization titers were created using the LANL Heatmap tool (https://www.hiv.lanl.gov/content/sequence/HEATMAP/heatmap_mainpage.html).

### Neutralizing antibody analysis

Positive responses were defined as a neutralization titer IC80 or IC50 below 10 μg per milliliter. Because multiple sequences were obtained for each mother, we ran all statistical comparisons twice per bnAb: once using the most resistant Env to each bnAb for each participant and once using the most sensitive. Proportions of viruses resistant to all bnAbs were compared between groups using two-sided 0.05-level Fisher exact tests.

Neutralization titers were compared using two-sided 0.05-level Wilcoxon rank sum tests. Response rate confidence intervals were calculated by the Wilson score method (44). Correlation was measured using Kendall’s τ and tested using a two-sided 0.05-level exact test.

### Signature Analysis

Sites in bnAb-specific contact regions where mutations are known to allow neutralization escape were identified using the LANL Env Feature Database (https://www.hiv.lanl.gov/components/sequence/HIV/featuredb/search/env_ab_search_pub.comp) under the feature types “contact”, “neutralization”, and “signature”, and only included in the analysis if more than one residue was observed at that position in the two env sets combined. Amino acid frequency rates at these sites were compared between groups using two-sided 0.05-level Barnard’s tests, and 95% confidence intervals were calculated by the Wilson score method (44).

Env characteristics (length, electrochemical charge and number of glycosylation sites) for the five variable regions were compared between BM and contemporaneous adult viruses using two-sided 0.05-level Wilcoxon rank sum tests. The electrochemical charge was calculated assigning −1 to amino acids D and E, and +1 to and amino acids H, K, and R. Logos were generated using the LANL tool AnalyzeAlign (www.hiv.lanl.gov/content/sequence/HIV/HIVTools.html). Analyses ran on R Studio, version 4.0.4, using packages tidyverse, officer, ggplot2, and ggpubr.

## Data Availability

All sequences were previously submitted to GenBank under accession numbers HM215361-HM215362m HM070481, HM070482, HM070484, HM070516, HM070517, HM070518, HM070525, HM070564, HM070569, HM070570, HM070604, HM070606, HM070612, HM070620, HM070717, HM070721, HM070724, HM070737, HM070749, HM070818, HM070822, HQ595836, HQ595839, HQ595847, HQ595852, FJ443316, FJ443382, FJ443533, FJ443575, FJ443639, FJ443670, FJ443711, FJ443808, FJ443841, FJ443999, FJ444017, FJ444059, FJ444103, FJ444124, FJ444586, HM215319, HM215324, HQ595742, HQ595747, HQ595748, HQ595749, HQ595750, HQ595757, HQ595759, HQ595763, HQ595766, HQ615942, HQ615943, HQ615944, HQ615945, HQ615946, HQ615950, HQ615951, HQ615952, HQ615954, HQ615955, HQ615959, JN681220-JN681223, JN681226-JN681234, JN681238-JN681249, JN681252-JN681256, JN967790, JN967792, JN967793, JN967794, JN967797, JN967798, JN977604, JN983803-JN983805, JQ061131, JQ352785, JQ352794, JQ352801, JX131327, KC154015, KC154015-KC154020, KC154022-KC154025, KF114881, KF114882, KF114884, KF114885, KF114889, KF114891, KF114893, KF114894, KF114895, KJ700458.

## Funding/Acknowledgements

This work was supported by NIH grants UM1AI068635 (HJ, KG), P01 AI117915 (SRP), and R01 AI162245 (SRP, EEG, and GF), and by a Collaboration for AIDS Vaccine Discovery (CAVD) grant (INV-007368 & INV-036842) from the Bill & Melinda Gates Foundation.

## Author Contributions

Manuscript conceptualization: EEG, HJ, DCM. Data curation: DCM, SRP, GF, ED. Statistical Analyses: EEG, KG, HJ. Figure preparation: KG, EEG. Manuscript writing: EEG, HJ. Manuscript editing: EEG, HJ, DCM, GF.

---

**Fig S1.**
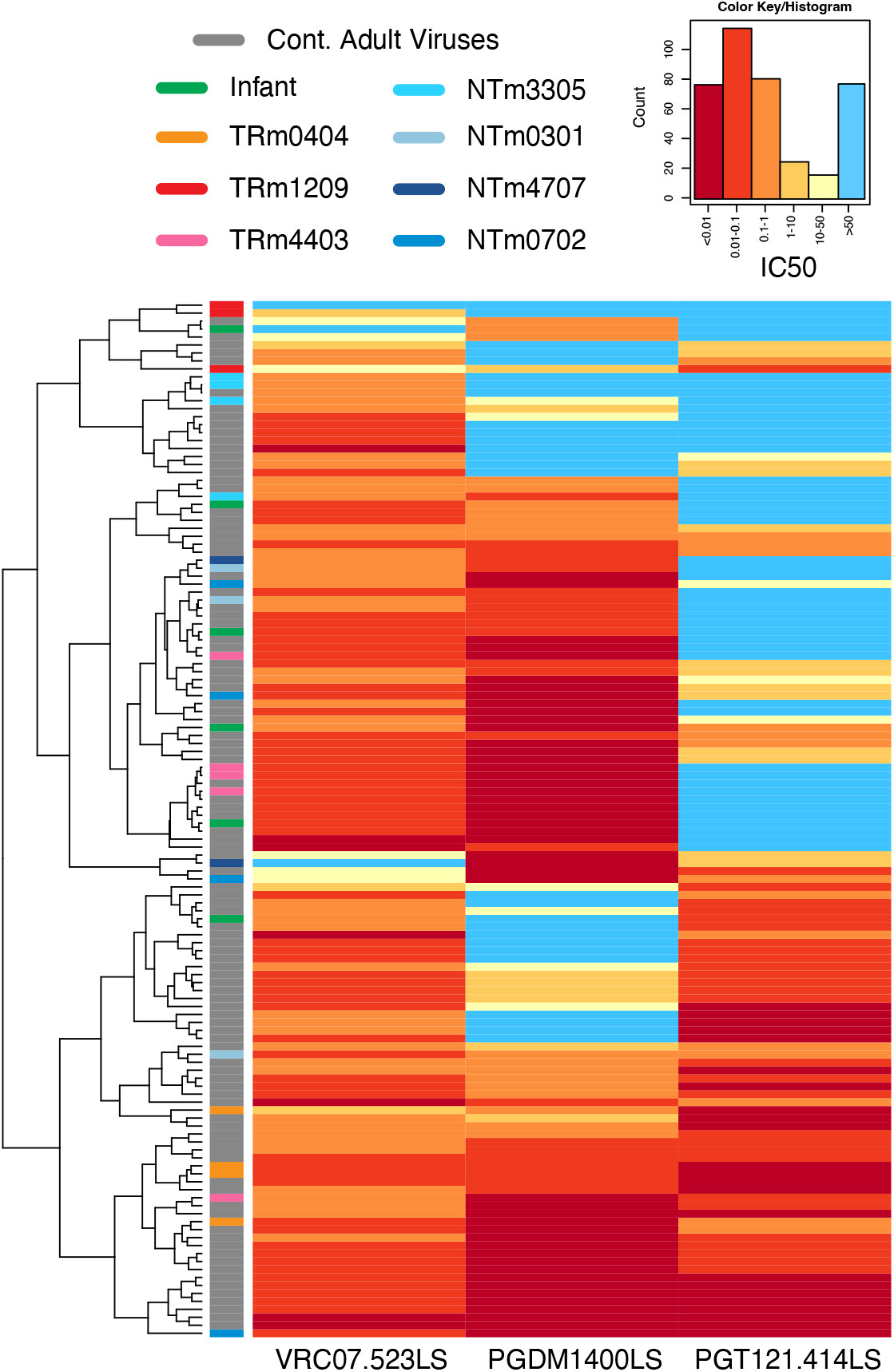
Heatmap visualization of the IC50 titers of all 130 Envs (rows) against the three bnAbs VRC07.523, PGT121, and PGDM1400 (columns). Titers are color-coded with warmer colors indicating stronger susceptibility and lighter colors indicating resistance, with light blue indicating above titers above the detection threshold of 50. Rows are hierarchically clustered and color-coded bars follow the same schema as in Fig. 1: warm colors for transmitting mothers, cold colors for non-transmitting mothers, and gray for contemporaneous adult viruses.

**Fig S2.**
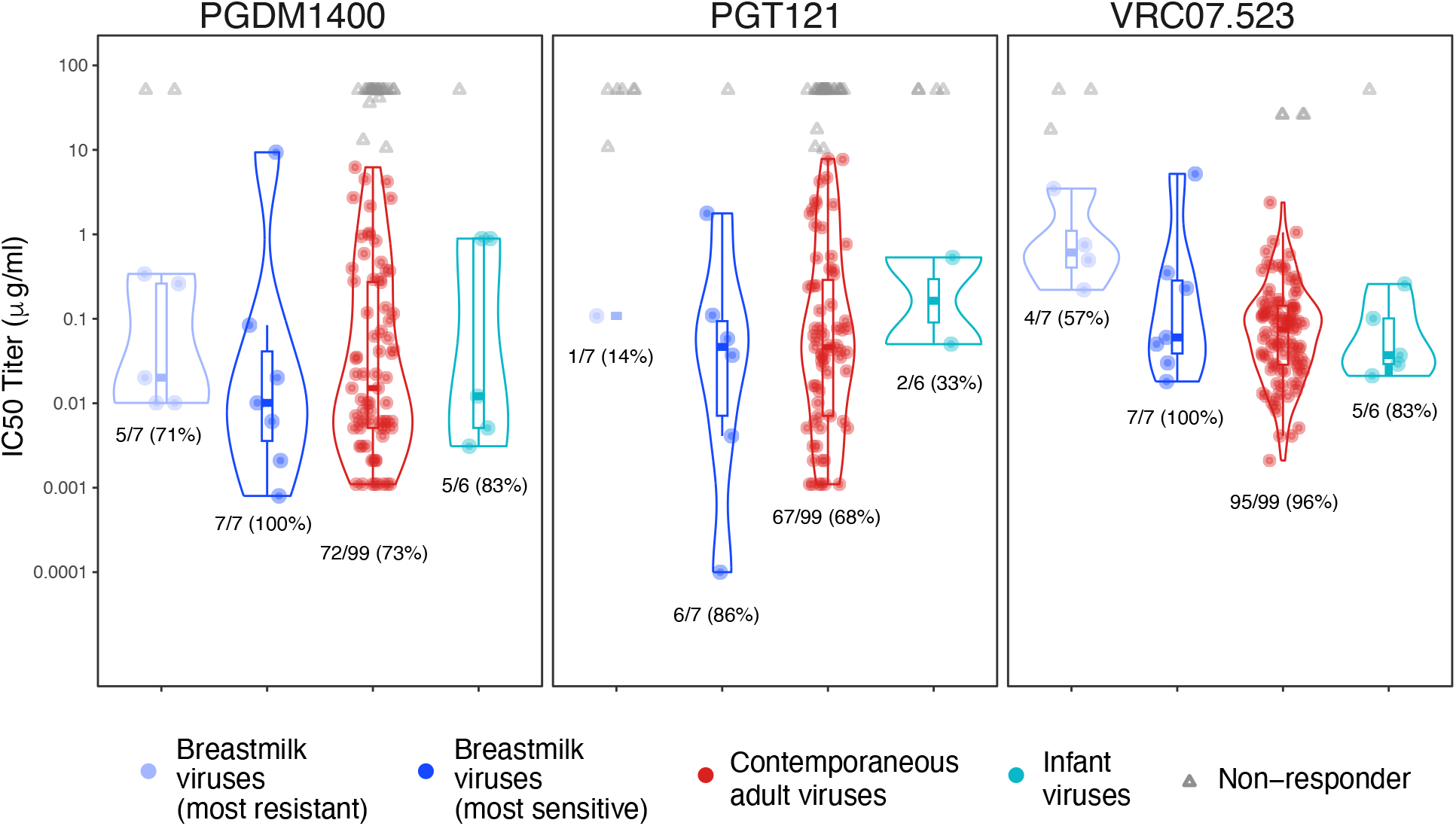
Distribution of IC50 titers against VRC07.523, PGT121, and PGDM1400, for breastmilk viruses, contemporaneous adult viruses, and infant viruses. Two sets of breastmilk viruses are considered: the virus most sensitive to a given bnAb, among all viruses sequenced from each mother; and the virus most resistant to a given bnAb.

**Fig S3.**
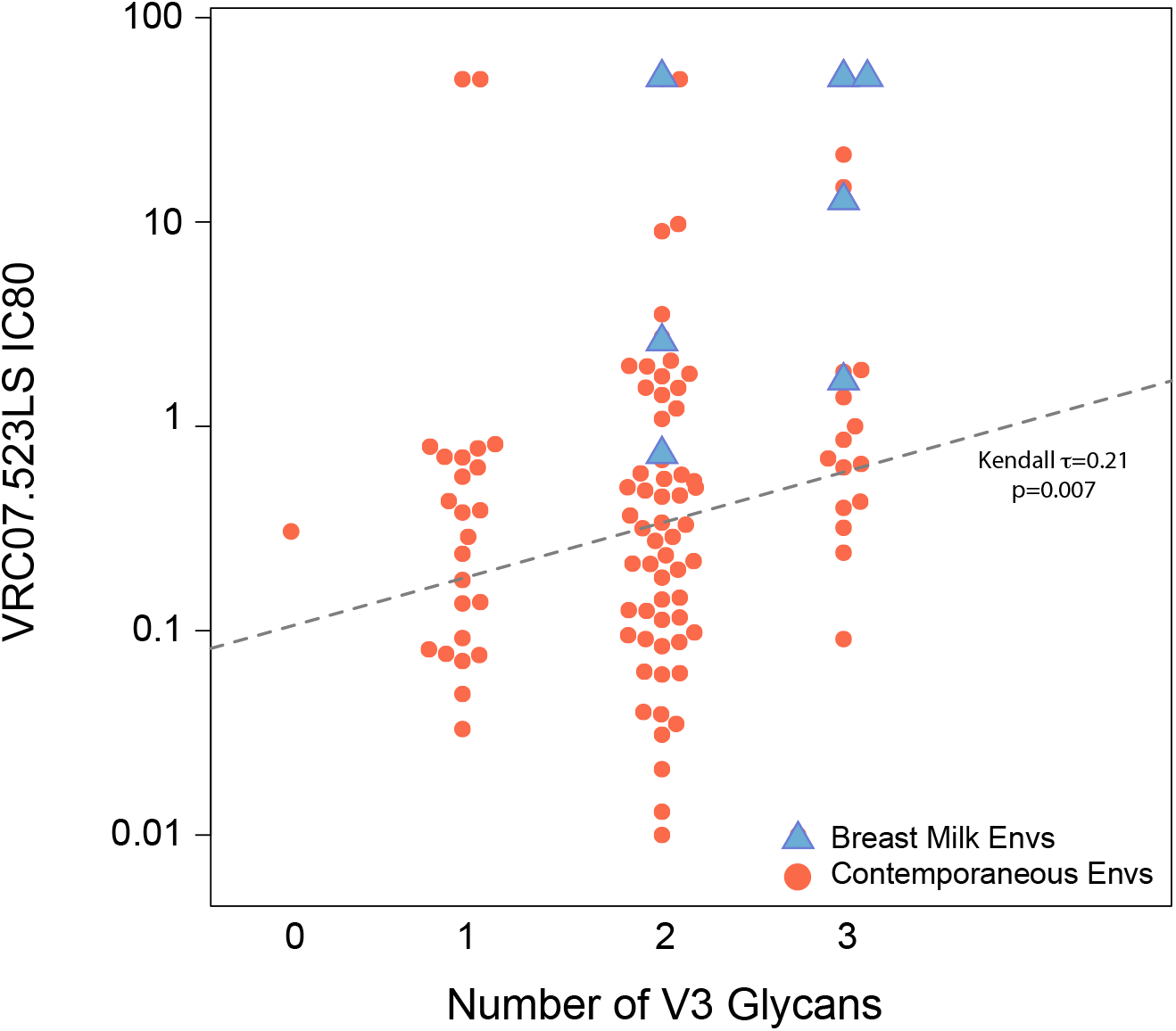
Correlation between the number of glycosylation sites in V3 and VRC07.523 resistance. The 7 VRC07.523 most resistant BM envs are shown in light blue triangles while the 99 contemporaneous clade C viruses are shown in orange circles.

**Fig S4.**
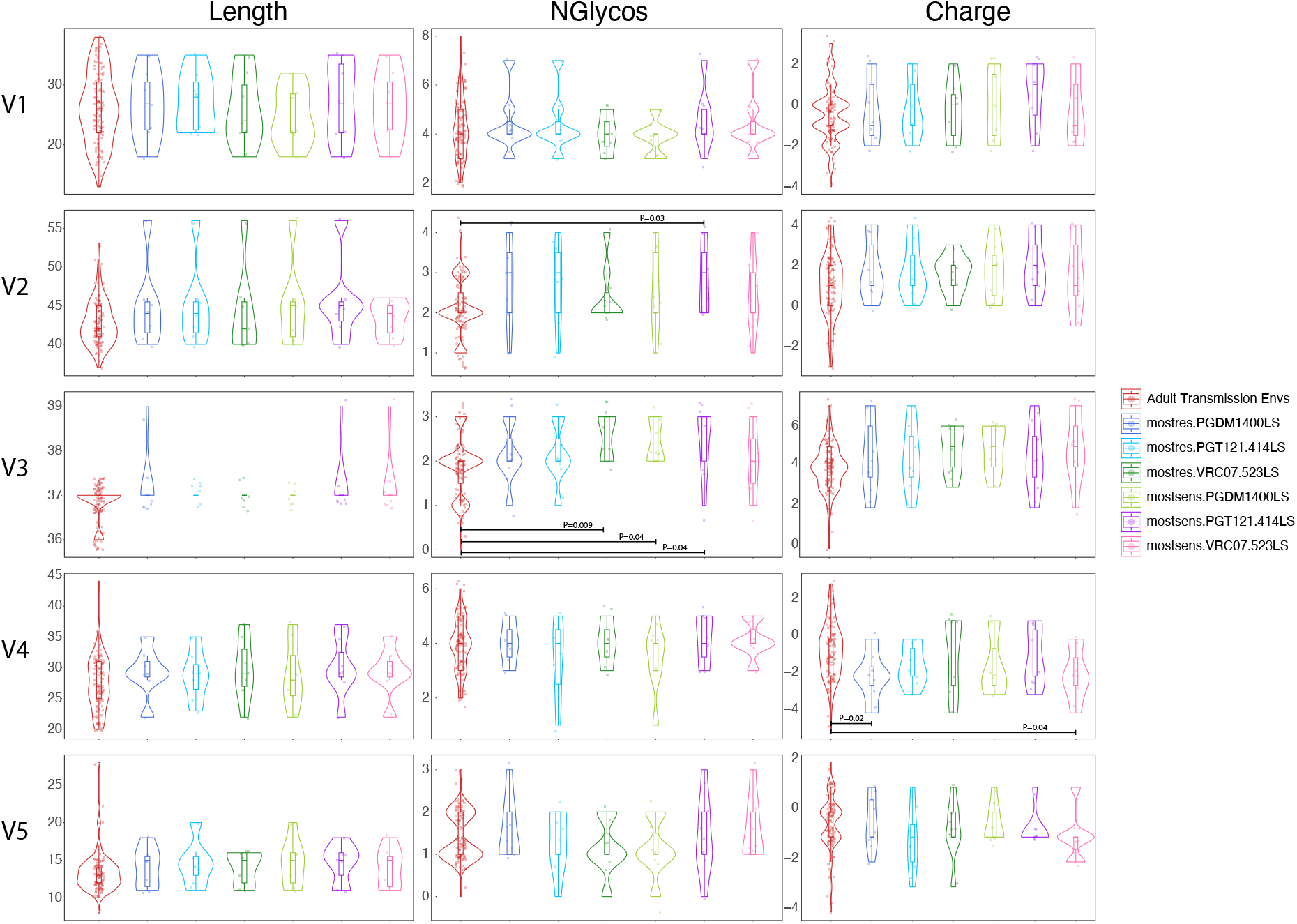
Comparisons of length, charge, and number of glycosylation sites at the five variable regions V1-V5. Each violin plot represents a different dataset. Statistically significant comparisons are marked with the corresponding p-value. All other comparisons were not statistically significant at a 0.05 significance threshold.

